# Bisphenol S moderately decreases the expression of syncytiotrophoblast marker genes and induces apoptosis in human trophoblast lineages

**DOI:** 10.1101/2023.10.19.563141

**Authors:** Enoch Appiah Adu-Gyamfi, Joudi Salamah, Elisha Ann Cheeran, Bum-Kyu Lee

## Abstract

Bisphenol S (BPS) is currently used in the manufacturing of several household equipment such as water pipes and food containers. Hence, its entrance into the human body is almost inevitable. The presence of BPS in body fluids has been reported. However, its potential toxicity, especially on human placenta development and pregnancy progression, has not been explored. In this study, we assessed the impacts of BPS on self-renewal and differentiation potentials of placental stem cells, also known as trophoblast stem cells (TSCs), by exposing them to three different BPS concentrations during both self-renewal and differentiation of TSCs into syncytiotrophoblast (ST), extravillous trophoblast (EVT), and trophoblast organoids. Interestingly, BPS treatment did not affect the stemness, cell cycle and proliferation of the TSCs but it induced apoptosis in each trophoblast lineage. BPS altered the expression of several fusion-related genes. However, this alteration did not translate into significant morphological defects in the STs and organoids. Moreover, BPS did not impair the differentiation of TSCs into EVTs. These findings suggest that the presence of BPS at the feto-maternal interface may exaggerate trophoblast apoptosis and moderately inhibit the trophoblast fusion pathway to affect placenta development and pregnancy. Our study offers valuable insights into the potential toxicity of BPS on human placenta development, emphasizing the need for epidemiological assessment of the relationship between maternal serum levels of BPS and pregnancy complications.

## 1. Introduction

The human placenta is a transient organ that is indispensable for pregnancy progression and maternal health. The major cell type in the placenta is the trophoblast, which has three main lineages: cytotrophoblasts (CTs), syncytiotrophoblasts (STs) and extravillous trophoblasts (EVTs). During placenta development, the CTs, which are stem cells, proliferate and differentiate into the STs and EVTs, with a continuous occurrence of strictly controlled apoptosis to remove superfluous and dysfunctional cells (Adu-Gyamfi et al., 2021; Knöfler et al., 2019). While the EVTs invade the uterus to establish the uteroplacental circulation and the histotrophic nutrition, the STs serve as a barrier that controls the exchange of substances between the fetus and the mother (Adu-Gyamfi et al., 2021; Knöfler et al., 2019). It has been found that abnormal proliferation, differentiation, and apoptosis of the trophoblasts lead to defective placenta and pregnancy complications such as miscarriage, preeclampsia, and fetal growth restriction (Adu-Gyamfi et al., 2020b; Adu-Gyamfi et al., 2019; Erel et al., 2001; Farah et al., 2020; He et al., 2013; Sharp et al., 2010; Zhang et al., 2020). Accumulation of xenobiotics at the feto-maternal interface is believed to be among the contributing factors to these anomalies.

Some xenobiotics can compromise or exaggerate the functionality of several cells, tissues, and organs. One group of these chemicals is the bisphenols, among which are bisphenol A (BPA) and bisphenol S (BPS). BPA is used in the production of food and beverage containers, water pipes, medical equipment, coatings, adhesives, high-performance composites, and automotive and aircraft parts (Adu-Gyamfi et al., 2022). Ever since it was shown that BPA significantly affects several developmental processes, such as impairing decidualization (Nelson et al., 2020), inducing abnormal placentation, and causing pregnancy complications (Adu-Gyamfi et al., 2022), its use has been banned in some geographical settings. BPS has now been recommended as an alternative substitute for BPA. Hence, this chemical is currently used in the production of several equipment which were previously produced with BPA. Due to this, human exposure to this chemical is now on the rise. Contemporary evidence suggests that BPS is detectable in the follicular fluid (Žalmanová et al., 2023), blood, and urine (Jin et al., 2018; Smarr et al., 2018), raising concerns about its potential toxicity. Therefore, it is imperative to conduct a comprehensive examination of the potential toxic effects of this chemical to ascertain whether it is safe to the human body and pregnancy. It has been demonstrated that BPS affects EGF signaling in human trophoblast cells (Ticiani et al., 2022), impairs endocrine functions in the sheep placenta (Gingrich et al., 2018), and induces morphological defects in the junctional zone of the mouse placenta (Mao et al., 2020). However, the impacts of BPS on human placenta development remains to be clarified.

Previous research about the effects of xenobiotics on placenta development primarily relied on animal models and immortalized or choriocarcinoma trophoblast cell lines such as JEG3, Bewo and HTR8/SVneo cells. These models exhibit varying gene expression signatures and do not closely mirror human placenta development. Therefore, experimental findings from them are not directly applicable to humans (Abbas et al., 2020; Hemberger et al., 2020). Since it is unethical to administer xenobiotics to pregnant women, it would be ideal to use an in vitro model that closely mirrors the human placental micro-environment for studying the toxic effects of xenobiotics on placenta development. The use of human trophoblast stem cells (TSCs) has been recommended in that regard (Adu-Gyamfi et al., 2022). The TSCs have the ability to differentiate into trophoblast lineages that have morphological characteristics and transcriptomic signatures which closely resemble those of the early human placenta (Okae et al., 2018). Therefore, they serve as a better model for studying the mechanisms of early human placenta development.

In this study, we aimed to determine the impacts of BPS on trophoblast stemness, cell cycle, proliferation, differentiation, and apoptosis. We differentiated human TSCs (which are the in vitro counterparts of the CTs) into STs, EVTs, and organoids in the presence or absence of BPS and then assessed the effects. Our findings will contribute to policy formulation regarding BPS use.

## 2. Materials and methods

### 2.1 TSC culture

Human TSCs were grown in TSC culture medium [Dulbecco’s modified Eagle’s medium (DMEM)/F12 (Gibco), New York, USA] supplemented with 0.2% fetal bovine serum (FBS; GeminiBio, California, USA), 1% insulin-transferrin-selenium-ethanolamine (ITS-X; Gibco), 0.1 mM β-mercaptoethanol (Sigma-Aldrich, St Louis, Washington, USA), 0.3% bovine serum albumin (BSA; GeminiBio), 0.5% penicillin– streptomycin (Gibco), 0.5 μM A83-01 (Wako Pure Chemical Corporation, Osaka, Japan), 0.5 μM CHIR99021 (Selleck Chemicals, Texas, USA), 0.5 μM SB431542 (Stemcell Technologies, Washington, USA), 0.8 mM valproic acid (VPA; Wako Pure Chemical Corporation), 5 μM Y27632 (ROCK inhibitor, Selleck Chemicals), 1.5 μg/ml L-ascorbic acid (Sigma-Aldrich) and 50 ng/ml epidermal growth factor (EGF; PeproTech, Cranbury, New Jersey, USA) in accordance with a previous protocol (Okae et al., 2018). Cell culture plates that had been coated with 5 μg/ml of Collagen IV (Corning, Bedford, Massachusetts, USA) were used for the cell culture. On reaching 70–80% confluency, the cells were detached with TrypLE (Gibco) for 8 min at 37°C and then passaged at a 1:3 split ratio. TSCs were cultured in a 37°C and 5% CO_2_ incubator. The medium was replaced every day with a fresh medium.

### 2.2 Differentiation of TSCs to STs

Previously cultured TSCs at ∼80% confluence were trypsinized. 3 × 10^5^ of the TSCs were seeded in 6 cm Petri dishes and cultured in 3 mL of ST medium [DMEM/F12 supplemented with 2 μM forskolin, 0.3% BSA, 1% ITS-X supplement, 0.1 mM 2-mercaptoethanol, 4% KSR, 50 ng/ml EGF, 0.5% Penicillin-Streptomycin, and 2.5 μM Y27632] in accordance with a previous protocol (Okae et al., 2018). On the second and fourth days, the medium was replaced with a fresh medium. On the 5th day, the STs were collected, and washed to remove dead cells after which RNA and proteins were extracted from them.

### 2.3 Differentiation of TSCs to EVTs

Trypsinized TSCs were seeded in 6-well plates pre-coated with 1 μg/ml of Collagen IV at a density of 1 × 10^5^ cells per well and grown in 2 mL of EVT medium [DMEM/F12 supplemented with 0.3% BSA, 0.1 mM 2-mercaptoethanol, 4% KSR, 0.5% Penicillin-Streptomycin, 1% ITS-X supplement, 2.5 μM Y27632, 100 ng/ml NRG1, and 7.5 μM A83-01] in accordance with a previous protocol (Okae et al., 2018). Shortly after suspending the cells in the medium, Matrigel (Corning, Bedford, Massachusetts, USA) was added to a final concentration of 2%. On day 3, the medium was replaced with EVT medium without NRG1, and Matrigel was added to a final concentration of 0.5%. On day 6, the medium was replaced with EVT medium without NRG1 and KSR. Matrigel was then added to a final concentration of 0.5%. The cells were harvested on day 8 for further experiments.

### 2.4 Differentiation of TSCs to organoids

TSCs at ∼80% confluence were trysinized. 5 x 10^3^ TSCs were suspended in 50 µL of 72% Matrigel droplet in Advanced DMEM/F12 medium. 5 droplets of the cell-matrigel mixture were introduced into the middle of 6 cm Petri dishes. It was ensured that the droplets were separated from one another and never adhered to the walls of the Petri dishes. The droplets were polymerized by putting the Petri dishes in a humidified 5% CO_2_ incubator at 37 ^O^C for 30 mins. The Petri dishes were then removed from the incubator, and then 2ml of trophoblast organoid medium was added to the droplets and the Petri dishes put back in the incubator. The medium was replaced every 2 days. The culture was continuously monitored for the appearance of small organoid clusters. On day 8, the organoids were harvested, and total RNA and proteins were extracted from them. The trophoblast organoid medium comprised Advanced DMEM/F12 supplemented with N2, B27, 2 mM L-glutamine, 100 ng/mL Recombinant human FGF-2, 500 nM A83-01, 1.5 µM CHIR99021, 50 ng/mL Recombinant human EGF, 1.25 mM N-Acetyl-L-cysteine, 50 ng/mL Recombinant human HGF, 100 μg/mL Primocin, 80 ng/mL Recombinant human R-spondin-1, 2.5 µM Prostaglandin E2, and 2 µM Y-27632, same as the medium that was used in a previous study in which organoids were established from human primary trophoblast cells (Turco et al., 2018).

### 2.5 Treatment of the TSCs, STs, EVTs, and organoids with BPS

BPS (Sigma-Aldrich, CAS-No:80-09-1, Purity: 98%) was dissolved in absolute ethanol, and then transferred into cell culture media to yield final concentrations of 0.01, 0.1 and 1 µM, which are concentrations that lie within the reported urine and blood levels of BPS in humans (Hehn, 2016; Jin et al., 2018; Smarr et al., 2018). Since TSCs, STs, EVTs, and organoids grow and develop in special media, specific BPS-containing media and their negative controls (NCs) were prepared for each model. The TSCs were treated with the various BPS concentrations for 2 days and were then harvested for cell cycle analysis, apoptosis assessment, and total RNA extraction. Differentiation of the TSCs into STs, EVTs, and organoids was done as already described above, with the only modification being the addition of BPS and ethanol to the media. On day 5, the STs were harvested, and on day 8, the EVTs and organoids were harvested for subsequent experiments.

### 2.6 Cell proliferation assay

The CCK8 kit (Abcam, Cambridge, Massachusetts, USA) was used to assess the proliferative rate of the TSCs. 3 × 10^3^ cells were seeded in 96-well plates in triplicates. After treating the TSCs with 0.01, 0.1 and 1µM of BPS and the NCs for 1 day, 2 days or 3 days in 100ul of media per well, we added 10 μl of the CCK8 solution to each well, and then incubated the cells for 1 h at 37 ◦C. We measured the optical density at 450 nm with a microplate reader (Thermo Fisher Scientific, MA, USA).

### 2.7 Flow cytometric analysis of the cell cycle and apoptosis

Only TSCs were used for cell cycle analysis while both TSCs and EVTs were used for the apoptosis assay. The cells were trypsinized and washed. For cell cycle analysis, about 1×10^5^ TSCs were fixed with 70% ethanol and incubated for, at least, 3 h at −20°C. After removing the ethanol, 150ul of Muse Cell Cycle reagent (Millipore, MCH100106) was used to incubate the cells for 30 mis at room temperature in the dark. The cells were analyzed by using the Muse flow cytometer. For apoptosis assay, about 1×10^5^ TSCs or EVTs were incubated with 150ul of Muse Annexin V & Dead Cell reagent (Millipore, MCH100105) for 20 mins at room temperature in the dark. The cells were then analyzed with the Muse flow cytometer (Luminex, Watertown, Massachusetts, USA).

### 2.8 Total RNA extraction and RT-qPCR

Total RNA was extracted from the TSCs, STs, EVTs and the organoids by using the single cell RNA purification kit (Norgen Biotek Corp, 51800, Ontario, Canada). The quality of the RNA was evaluated with 1% agarose gel electrophoresis and UV spectrophotometry. Complementary DNAs (cDNAs) were generated from 300 ng of total RNA using the SuperScript^TM^ III first strand synthesis SuperMix kit (Thermo Fisher Scientific, 18080400) and then diluted 20 times with DNase-free water. RT-qPCR was carried out by using the PowerTrack^TM^ SYBR Green Master Mix (Thermo Fisher Scientific, A46110). Primers were designed to amplify the junction between two exons using a web-based primer design program, Primer 3 (http://bioinfo.ut.ee/primer3/). The primer sequences are listed in Supplementary Table 1. Cycle threshold (Ct) values of the samples and the controls were normalized against GAPDH while the relative gene expression was calculated as fold enrichment by using the 2^−ΔΔCT^ method. The relative gene expression was calculated from an average of at least two replicates.

### 2.9 Protein extraction and Western blot

4X Laemmli sample buffer (Bio-Rad, USA) supplemented with 2-Mercaptoethanol was added to the STs or organoids, and the mixture homogenized. The mixtures were incubated at 90 ◦C for 10 min, after which appropriate concentrations of SDS-PAGE gel were prepared and used for electrophoresis. The proteins in the gels were transferred to PVDF membranes (Bio-Rad, USA). The non-specific binding sites on the membranes were blocked with 5% non-fat skim milk (Foodhold, USA) at 25 ◦C for 60 min. The membranes were then probed with primary antibodies at 4 ◦C for 16h. The primary antibodies were specific for SDC1, CGB3, CSH1, CDH1 and GAPDH (Santa Cruz, USA). For each primary antibody, the concentration used was 1 μg/ml. GAPDH was used as a control to ensure that protein loading was equal across the wells in the gels. After washing the membranes three times with PBST, we incubated them with HRP-conjugated goat anti-mouse IgG (0.5 μg/ml, Active Motif, USA) at 25 ◦C for 1 h. After this, the membranes were washed three times with PBST. Protein bands were then detected in the membranes by using the ECL chromogenic solution (Thermo Fisher Scientific, USA). The protein bands were quantified by using the ImageJ software.

### 2.10 Statistical analyses

The data were analyzed by using the GraphPad Prism 9 software. Error bars in the figures indicate mean ± SD (standard deviation) of, at least, two independent experiments. Significant differences between groups were assessed with two-tailed t-test. *, **, and *** denote significance levels of *P* < 0.05, P < 0.01, and P < 0.001, respectively.

## 3. Results

### 3.1. Successful establishment of the ST, EVT, and organoid models

We established in vitro models of the villous and extravillous trophoblasts of the human placenta by differentiating TSCs into STs, EVTs, and organoids (**Figs. 1A**). Each trophoblast lineage had morphological and gene expression characteristics that are comparable to those of the actual placenta. In the ST model, we verified the upregulation of ST markers, including fusogenic genes (*CGA, CGB3, CSH1, ERVW1, ERVFRD, GCM1, PSG1, SDC1,* and *CYP19A1*) (**Figs. 1B, 1C**, and **1D**) as well as downregulation of *TEAD4* (a stem cell marker) and *CDH1* (an epithelial marker) (**Fig. 1E**) compared to TSCs. In the EVT model, we confirmed the upregulation of EVT markers (*HLA-G, MMP2, ASCL2, ITGA5,* and *FN1*) (**Fig. 1F**) and a decrease in *TEAD4* expression (**Fig. 1G**) compared to TSCs. In the organoid model, we observed that the sizes of the organoids gradually increased during their 12 days formation period (**Fig. 1H**). Interestingly, the expressions of *CGA, CGB3, GCM1, PSG1, SDC1, CSH1,* and *CYP19A1* increased gradually and reached their maximal levels on day 8 of the organoid formation process and then declined afterwards (**Fig 1I**), suggesting that the organoids probably reached their maximal maturation on day 8. Hence, day 8 organoids were selected for further experiments. We further validated an increase in the expressions of *CGA, CGB3, CSH1, ERVW1, ERVFRD, GCM1, PSG1, SDC1,* and *CYP19A1* in the day 8 organoids compared to the TSCs (**Figs. 1J, 1K**, and **1L**). Notably, while there was a reduction in the expression of *CDH1*, there were no significant changes in the expressions of stem cell markers (*TEAD4, ELF5, GATA3*, and *TP63*) (**Fig. 1M**) and proliferation markers (*Ki67* and *PCNA*) (**Fig. 1N**). This is likely due to the organoids comprising a mixture of TSCs and STs. In summary, we successfully established in vitro models representing both villous and extravillous trophoblasts of the human placenta.

**Fig. 1.**
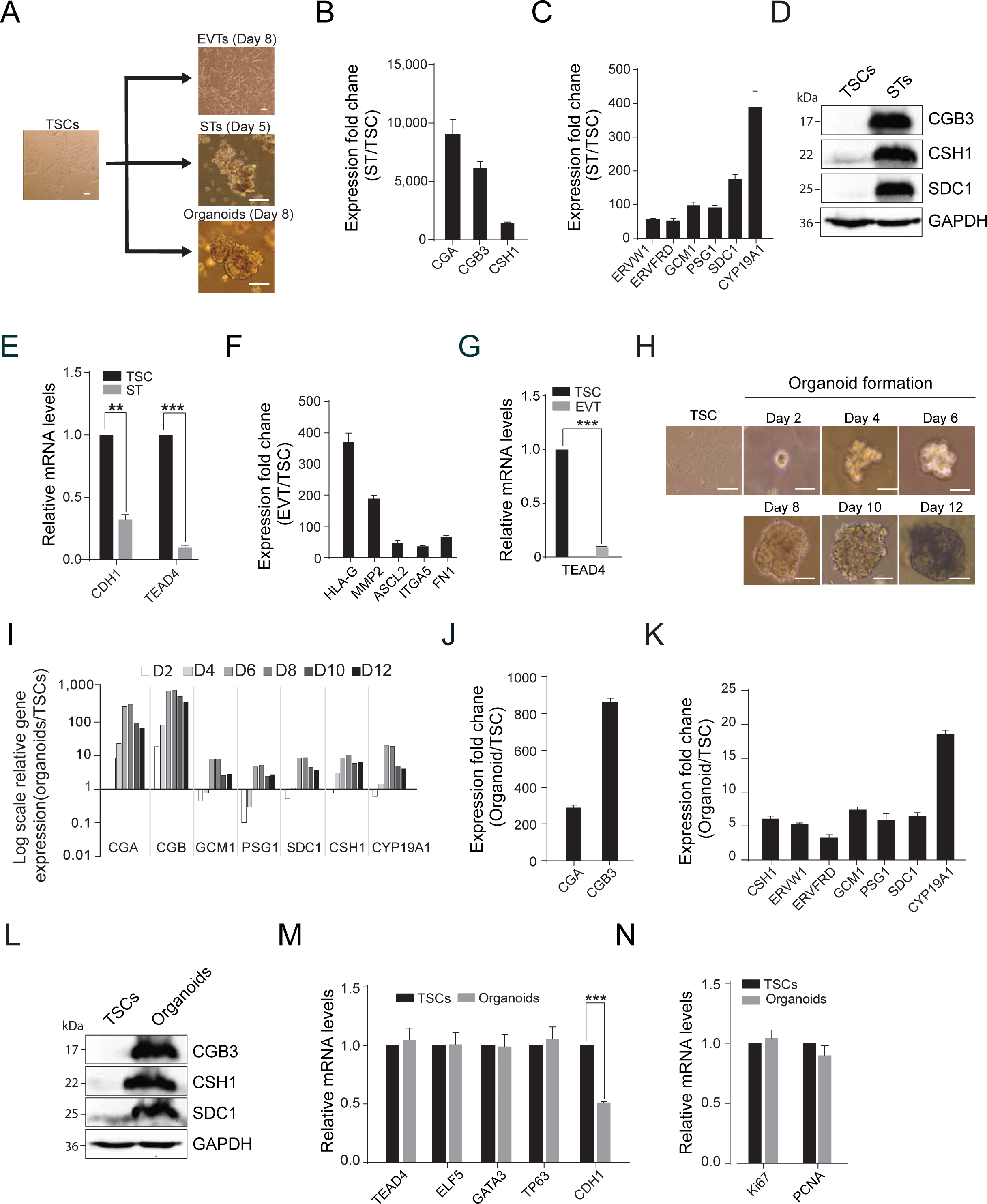
The established placental models and their characteristics. (**A**) Images showing the differentiation of TSCs into EVT, ST, and organoid models; (**B** and **C**) Expression fold changes of ST marker genes in STs relative to TSCs; (**D**) Protein levels of ST marker genes in TSCs and STs; (**E**) Transcript levels of epithelial (*CDH1*) and stem cell (*TEAD4*) markers in STs relative to TSCs; (**F**) Expression fold changes of EVT marker genes in EVTs relative to TSCs; (**G**) Transcript level of a stem cell marker, *TEAD4*, in EVTs relative to TSCs; (**H**) Images showing the time-course differentiation of TSCs into organoids; (**I**) Expression fold changes of ST marker genes during the time-course differentiation of TSCs into organoids; (**J** and **K**) Expression fold changes of ST marker genes in organoids relative to TSCs; (**L**) Protein levels of ST marker genes in TSCs and organoids; (**M** and **N**) Transcript levels of stem cell markers (**M**) and proliferation markers (**N**) in organoids relative to TSCs.

### 3.2. BPS elicits no significant effect on stemness, proliferation, and cell cycle of TSCs

We exposed TSCs to three different BPS concentrations (0.01, 0.1, and 1 μM) for 2 days, as illustrated in **Fig. 2A**. We observed that BPS treatment did not affect the expressions of stem cell marker genes (**Fig. 2B**). Consistently, proliferation assay showed no difference in growth rates between the BPS-treated and the NC TSCs (**Fig. 2C**), along with no significant expression change in proliferation marker genes (**Fig. 2D**). Flow cytometric analysis further indicated that the cell cycle did not differ between the BPS-treated and the NC TSCs (**Fig. 2E** and **Supplementary Fig. 1**). We also assessed the impact of each BPS concentration on the proliferation and stemness of the TSCs within the organoids. On day 8 of TSC differentiation into the organoids, we found that BPS treatment did not affect the expressions of the stem cell markers as well as the cell proliferation markers within the organoids (**Figs. 2F** and **2G**). Collectively, these results indicate that BPS has no significant effect on the self-renewal of TSCs.

**Fig. 2.**
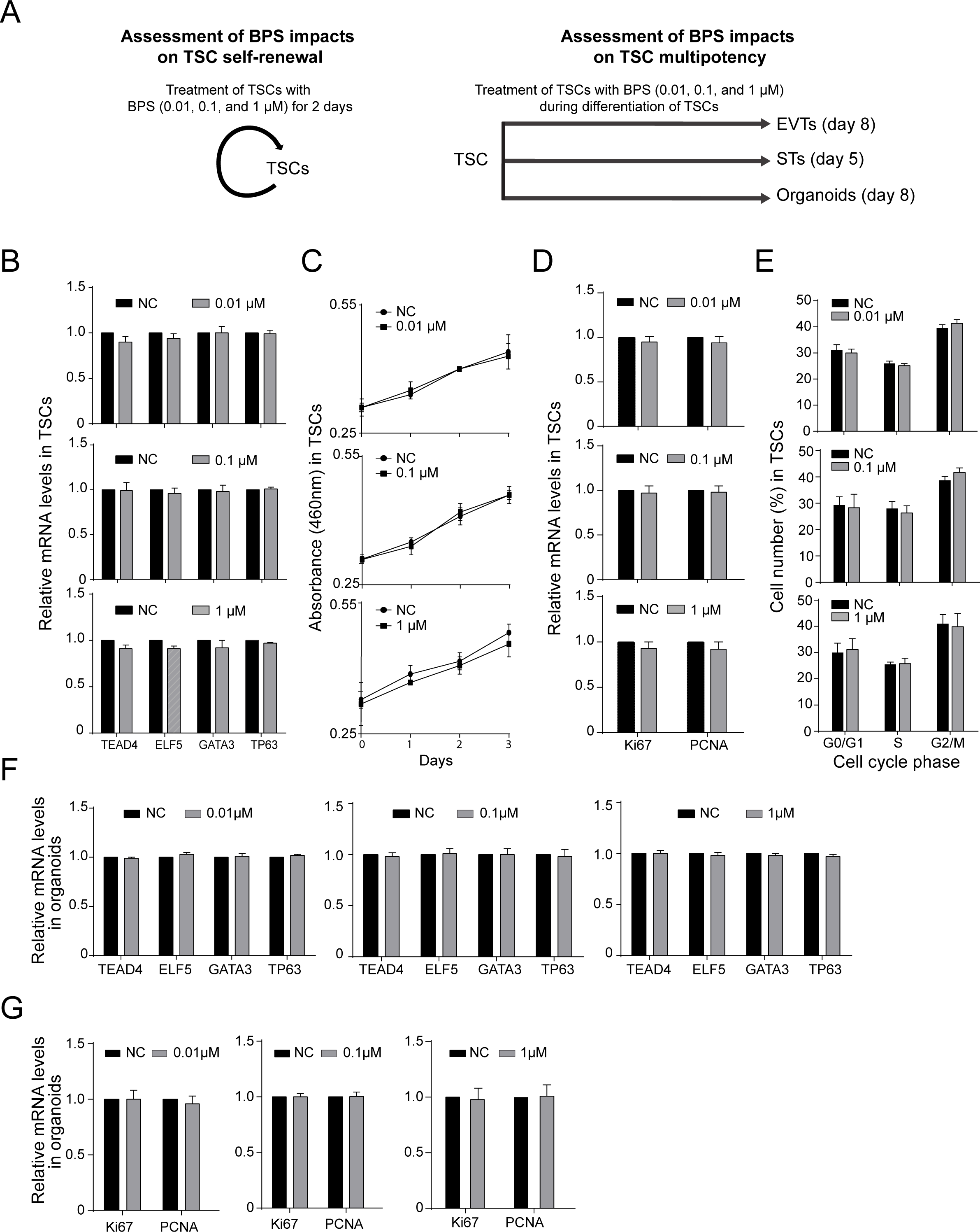
BPS does not affect TSC marker gene expression, cell cycle, and proliferation. (**A**) A schematic diagram illustrating the experimental design we used to investigate the impacts of BPS on the self-renewal of TSCs and their differentiation potentials. We treated TSCs with BPS for 2 days. Also, during the differentiation of TSCs into STs, EVTs, and organoids, we treated them with BPS (0.01, 0.1, or 1 μM). (**B**) Transcript levels of stem cell markers in the BPS-treated TSCs relative to NCs; (**C**) Proliferation rates in the BPS-treated and NC TSC; (**D**) Transcript levels of proliferation markers in the BPS-treated TSCs relative to NCs; (**E**) Cell population distribution in different cell cycle phases in BPS-treated and NC TSCs. (**F**) Transcript levels of stem cell markers in the BPS-treated organoids relative to NCs; (**G**) Transcript levels of proliferation markers in the BPS-treated organoids relative to NCs.

### 3.3. BPS moderately decreases the expression of ST marker genes in STs and organoids, while having no discernible effect on EVT formation

To explore the impacts of BPS on the multipotency of TSCs, we treated the TSCs with three different BPS concentrations (0.01, 0.1, and 1 μM) during differentiation of TSCs into STs, EVTs, and organoids (**Figs. 3A** and **4B**). On day 5 of ST formation, we observed no significant morphological differences between the BPS-treated STs and their corresponding NC STs (**Fig. 3A**). However, BPS treatment moderately reduced the expressions of several ST marker genes. For instance, in the STs treated with 0.01 μM of BPS, *CGB3, CSH1, GCM1,* and *PSG1* were downregulated while *CDH1* and *TEAD4* were upregulated. However, the expressions of *CGA, ERVW1, ERVFRD,* and *CYP19A1* remained unchanged compared to the NCs (**Fig. 3B**). Importantly, in the STs treated with 0.1 and 1 μM of BPS, we observed that 8 out of 9 ST marker genes (*CGA, CGB3, CSH1, ERVW1, GCM1, PSG1, SDC1* and *CYP19A1*) were downregulated while *CDH1* and *TEAD4* were upregulated compared to the NCs (**Fig. 3B**). This suggests that higher concentrations of BPS have stronger impacts on the downregulation of ST marker genes. Consistent with this, 1 μM of BPS moderately decreased the protein levels of CGB3, CSH1, and SDC1 while increasing the protein levels of CDH1 (**Fig. 3C** and **Fig. 3D**). Just like the STs, the organoids treated with BPS showed no significant change in morphology (**Fig. 3E**). However, a reduction in multiple ST marker genes was evident in this group compared to the NCs. These genes include *CGA, CGB3, CSH1, GCM1, ERVFRD, SDC1,* and *CYP19A* (**Fig. 3F**). Decreased protein levels of SDC1 and increased protein levels of CDH1 were also found in the organoids treated with 1μM of BPS (**Fig. 3G** and **3H**). On day 8 of TSC differentiation into EVTs, the BPS-treated EVTs and their corresponding NCs did not show significant morphological differences or changes in the expression of EVT marker genes (**Figs. 4A** and **4B**). Taken together, these findings suggest that BPS moderately inhibits the induction of ST marker genes during ST formation, while eliciting no effect on EVT formation.

**Fig. 3.**
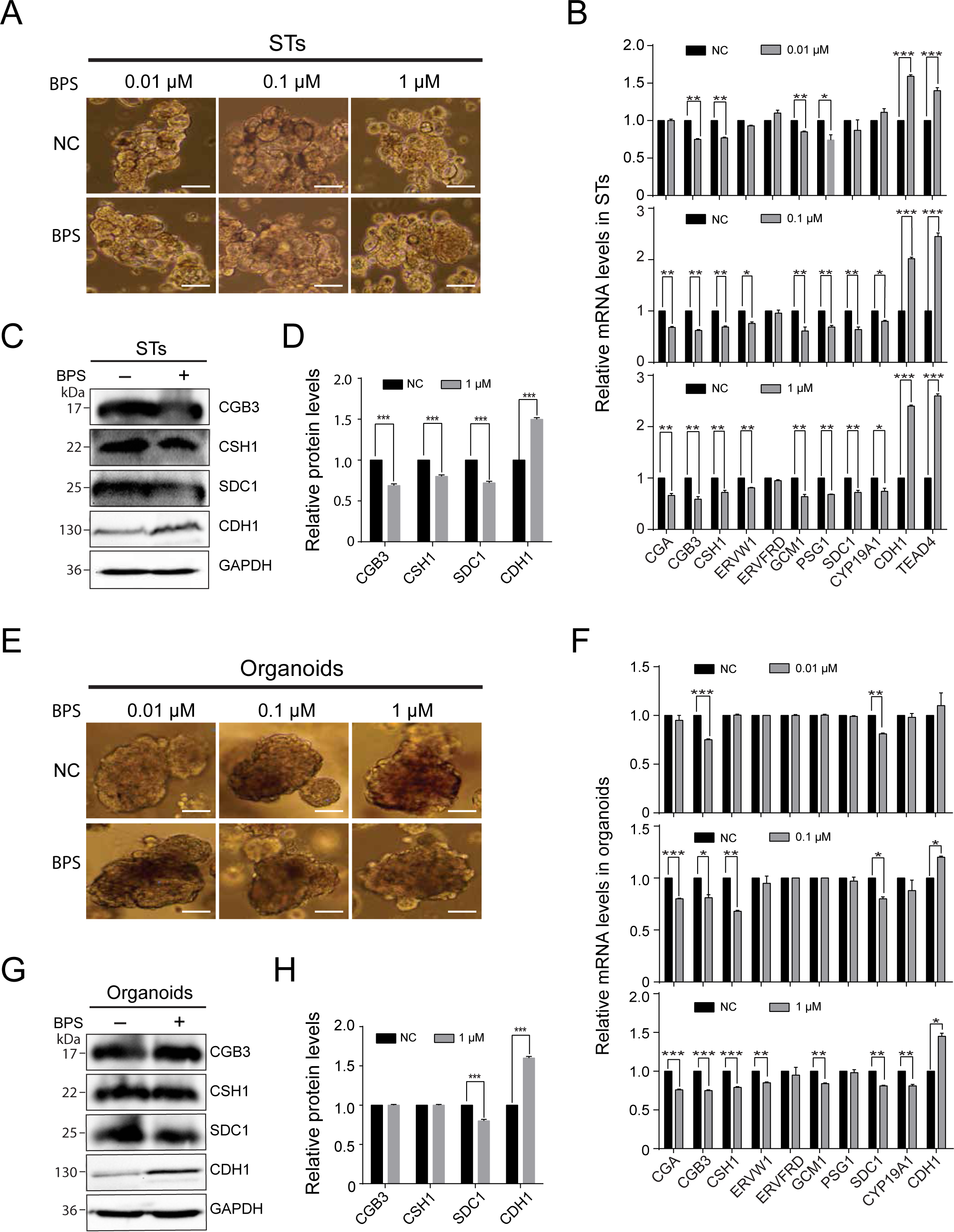
BPS moderately decreases the expression of trophoblast fusion marker genes in the STs and organoids. (**A**) Representative images showing BPS-treated and NC STs after 5 days of TSC differentiation into STs; (**B**) Transcript levels of trophoblast fusion, stem cell, and epithelial markers in the BPS-treated STs relative to NC STs; (**C**) Protein levels of ST marker genes in the BPS-treated STs and NCs; (**D**) Protein levels of ST marker genes in the BPS-treated STs relative to the NCs; (**E**) Representative images showing BPS-treated and NC organoids after 8 days of TSC differentiation into organoids; (**F**) Transcript levels of trophoblast fusion, stem cell, and epithelial markers in the BPS-treated organoids relative to NC organoids; (**G**) Protein levels of ST marker genes in the BPS-treated organoids and NCs; (**H**) Protein levels of ST marker genes in the BPS-treated organoids relative to the NCs

**Fig. 4.**
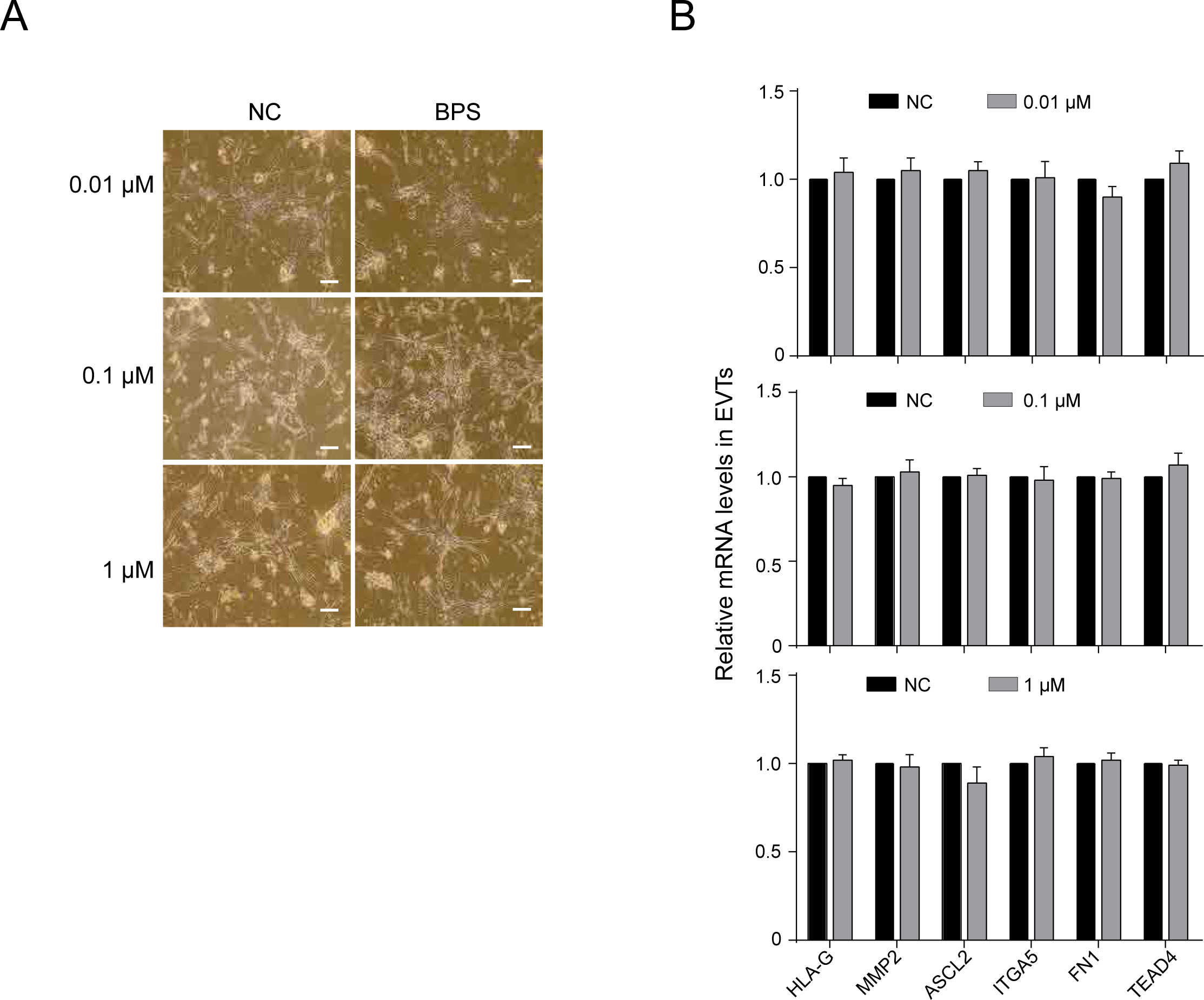
BPS does not affect EVT morphology and the expressions of EVT marker genes. (**A**) Representative images showing BPS-treated and NC EVTs after 8 days of TSC differentiation into EVTs; (**B**) Transcript levels of EVT markers in the BPS-treated EVTs relative to NC EVTs.

### 3.4. BPS induces apoptosis in trophoblast lineages

We determined the impacts of 0.01, 0.1 and 1 μM of BPS on the apoptosis of TSCs. Additionally, during the differentiation of TSCs into the STs, EVTs, and organoids, we assessed the impacts of these three different BPS concentrations on apoptosis in each trophoblast model. In the TSCs, we found that each BPS concentration increased the proportion of early and total apoptotic cells in a dose-dependent manner compared to the NC TSCs (**Fig. 5A** and **Supplementary Figs. 2** and **4A**). Additionally, we found an increase in the expressions of the pro-apoptotic genes (*BAX, TP53*, and *CDKN1B*) in the TSCs treated with each concentration of BPS (**Fig. 5B**). Treatment with BPS concentrations of 0.1 and 1 μM also resulted in a reduction in the expressions of the anti-apoptotic genes in the TSCs. Particularly, treatment with 1 μM of BPS significantly reduced the expression levels of *BCL2, SIRT1, MCL1, and PTMA* compared to the NC TSCs (**Fig. 5B**). In the EVTs, we found that only 1 μM of BPS induced apoptosis (**Fig. 5C** and **Supplementary Figs. 3** and 4B), leading to an increase in the proportion of late and total apoptotic cells compared to the NC EVTs. Further assessment showed that *BAX, TP53,* and *CDKN1B* were upregulated while *BCL2* was downregulated in the EVTs treated with 1 μM of BPS (**Fig. 5D**). However, unlike TSCs, the EVTs treated with 1μM of BPS did not show any change in the expressions of *SIRT1, MCL1,* and *PTMA* compared to the NC EVTs (**Fig. 5D**). In the ST model, 0.01 μM of BPS did not affect the expressions of the apoptosis markers. However, 0.1 and 1 μM of BPS were able to reduce the expressions of anti-apoptotic genes (*BCL2, SIRT1,* and *MCL1*) although they had no effect on the expressions of pro-apoptotic genes (**Fig. 5E**). In the organoid model, only the 1μM of BPS was able to downregulate *BCL2, SIRT1,* and *MCL1* while upregulating *TP53* and *CDKN1B*. However, it did not affect the expressions of *PTMA* and *BAX* (**Fig. 5F**). These findings suggest that BPS can induce apoptosis in trophoblast lineages in a dose-dependent manner.

**Fig. 5.**
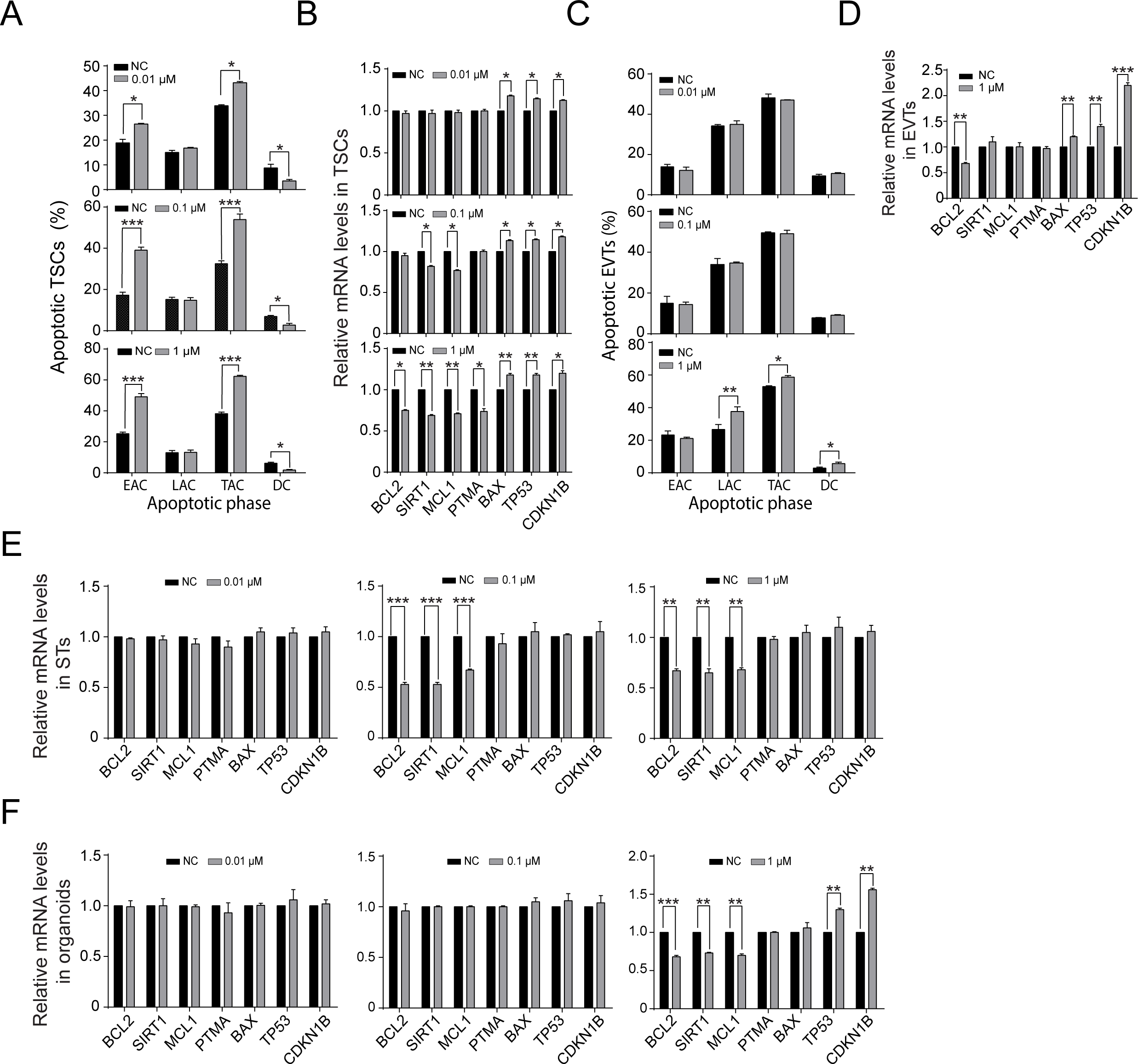
BPS induces apoptosis in TSCs, STs, EVTs, and organoids. (**A**) The distribution of cell population in different apoptotic phases (EAC: Early apoptotic cells, LAC: late apoptotic cells, TAC: Total apoptotic cells, and DC: Dead cells) of BPS-treated and NC TSCs; (**B**) Transcript levels of apoptosis markers in the BPS-treated TSCs relative to NC TSCs; (**C**) The distribution of cell population in different apoptotic phases of BPS-treated and NC EVTs; (**D**) Transcript levels of apoptosis markers in the 1uM BPS-treated and NC EVTs; (**E**) Transcript levels of apoptosis markers in the BPS-treated STs relative to NC STs; (**F**) Transcript levels of apoptosis markers in the BPS-treated organoids relative to NC organoids.

## 4. Discussion

In this study, we successfully established in vitro human placental models. In these models, the TSCs organized into either villous or extravillous structures that are similar to those of the actual placenta. Human placenta development mainly encompasses trophoblast proliferation, differentiation, and apoptosis which are critically regulated by a myriad of molecules, including transcription factors (Papuchova and Latos, 2022), hormones (Adu-Gyamfi et al., 2020c), and cytokines (Adu-Gyamfi et al., 2020a; Appiah Adu-Gyamfi et al., 2020). During differentiation, the CTs either transform into STs or EVTs. Formation of the STs, a process known as syncytialization, involves the fusion of the cell membranes of adjacent CTs, leading to protoplasmic merging (Kar et al., 2007). This occurrence is mediated by molecular processes such as upregulation of the syncytins (ERVW1, ERVFRD) (Roberts et al., 2021) and the hormonal genes (CGA, CGB3) (Okae et al., 2018) as well as the downregulation of epithelial markers such as CDH1 (E-cadherin) (Adu-Gyamfi et al., 2021). We confirmed these patterns of gene expression upon differentiation of TSCs into STs and organoids, providing evidence that we successfully established ST and trophoblast organoid models. Formation of the EVTs, a process known as epithelial-mesenchymal transition, is characterized by the CTs acquiring a spindle-shaped morphology and invasive abilities. This cellular process is mediated by the upregulation of HLA-G and mesenchymal markers such as ASCL2, MMP2, and ITGA5 (Okae et al., 2018; Varberg et al., 2021). As expected, we observed this expression pattern of the molecular markers in the EVT lineage we established. By using these models, we have demonstrated that BPS may moderately interfere with placenta development.

Proliferation of the CTs is the first stage in human placenta development. Both hypo- and hyper-proliferative rates of trophoblast cells are detrimental to the formation of a functional placenta. For instance, when CT proliferation is reduced, few CTs become available to undergo differentiation, leading to defective ST and EVT formation. Conversely, when proliferation is excessively stimulated, CT differentiation becomes inhibited, resulting in the impairment of ST and EVT formation. A previous study demonstrated that BPS does not affect the proliferation of HTR8/SVneo cells (Ticiani et al., 2022). This cell line is immortalized and has characteristics of the non-proliferative extravillous trophoblast cells which are completely different from the progenitor CTs or TSCs. By using TSCs and organoid models, we have shown that BPS does not affect the proliferation, cell cycle, and stemness of TSCs. Hence, BPS may not be detrimental to the proliferation and renewal of the trophoblast progenitor cells within the placenta.

At the feto-maternal interface, physiologic regulation of trophoblast differentiation is vital to the establishment of a normal placental architecture while dysregulation of this process, either due to the abnormal expression of some genes or the presence of a xenobiotic, induces placental abnormalities to affect pregnancy (Gong et al., 2021; Mohamad et al., 2020; Rosenfeld, 2021; Wu et al., 2023). In this study, we found that BPS did not induce noticeable defects in EVT morphology and the expression of EVT marker genes. We also found that BPS did not significantly affect the morphologies of the STs and organoids. However, it moderately reduced the levels of several ST formation marker genes. Additionally, it augmented the expression of the epithelial marker, *CDH1*, and the stem cell marker, *TEAD4* (Saha et al., 2020) in STs. Notably, BPS elicited these effects in a dose-dependent manner, with the highest concentration causing the most marked effect. Consistent with our observation, BPS impaired the placental cell fusion pathway but it did not affect the morphology of the sheep placenta (Gingrich et al., 2018). Similarly, in a previous in vitro study, ∼0.8 μM of BPS did not affect the morphology of spontaneously fused primary CTs (Ticiani et al., 2021). Collectively, these findings suggest that BPS may cause a moderate molecular-level defect which does not translate into significant morphological defects in STs. Besides, BPS may have no significant effect on CT differentiation into EVTs during placenta development.

Trophoblast apoptosis is an integral process in placenta development. The interplay between pro- and anti-apoptotic regulators at the feto-maternal interface tightly regulates the rate of trophoblast apoptosis. During normal pregnancy, trophoblast apoptosis increases with advancing gestation. This serves to eliminate unwanted, superfluous, and dysfunctional cells; thereby, promoting placental growth and functionality. However, excessive apoptosis impairs placenta development to cause pregnancy pathologies (Raguema et al., 2020; Sharp et al., 2010; Straszewski-Chavez et al., 2005; Tomas et al., 2011). We have shown that BPS triggers apoptosis in each trophoblast lineage. This was evidenced by an increase in the proportion of total apoptotic TSCs and EVTs, an increase in the expression of pro-apoptotic genes and a reduction in the expression of anti-apoptotic genes in the TSCs, STs, EVTs, and the organoid model. While all the concentrations of BPS triggered apoptosis in the TSCs, only the higher BPS concentrations were able to induce apoptosis in the STs and EVTs. This suggests that CTs may be more susceptible to the apoptotic effects of BPS than STs and EVTs. Due to this apoptotic effect, the presence of BPS within the placental micro-environment may have a synergistic effect with physiological apoptosis; thereby, amplifying trophoblast apoptosis to the detriment of placenta formation and functioning.

## 5. Conclusions

This study is the first to use human TSCs and trophoblast differentiation models to assess the impacts of BPS on human placenta development. Our findings suggest that the presence of BPS at the feto-maternal interface may exaggerate trophoblast apoptosis while moderately inhibiting trophoblast fusion to impair placenta development. Since BPS and its products are in use, the global consumption of BPS is bound to rise exponentially in the coming decades. Hence, epidemiological studies should monitor BPS levels in the blood and placentas of pregnant women and assess the relationship between maternal serum levels of BPS and pregnancy complications. The findings will assist in policy formulations concerning the use of BPS and its products in daily life.

## Data Availability

The original data can be obtained from the corresponding author via an email.

## Acknowledgement

We are grateful to Dr. Takahiro Arima and Dr. Hiroaki Okae of Tohoku University for providing us with human TSCs and technical advice. This research was supported by start-up funding from the University at Albany-State University of New York

## Supplementary data

**Supplementary Fig.1.**
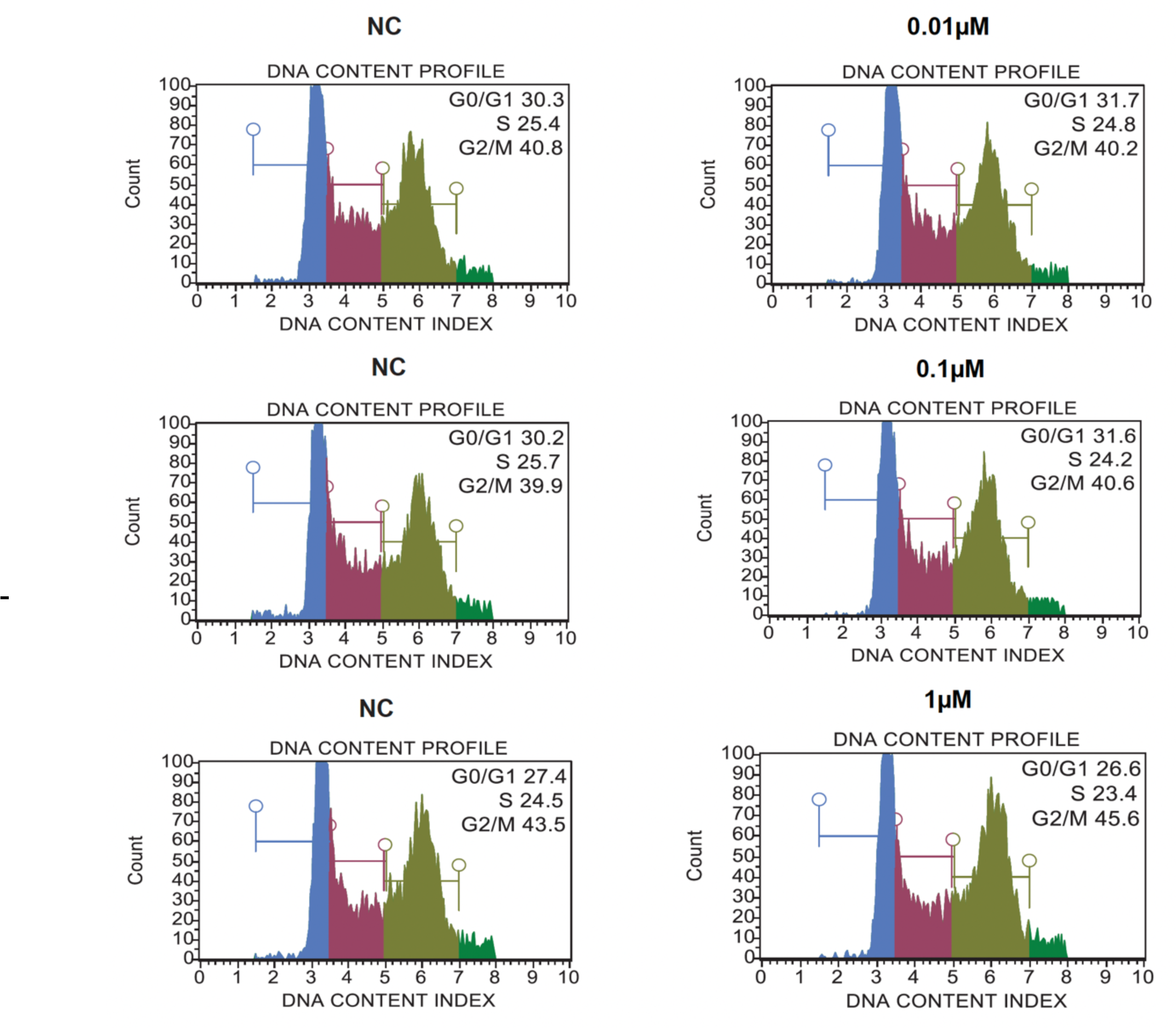
Representative histogram from cell cycle analysis presenting the proportion of cells in each phase of the cell cycle.

**Supplementary Fig.2.**
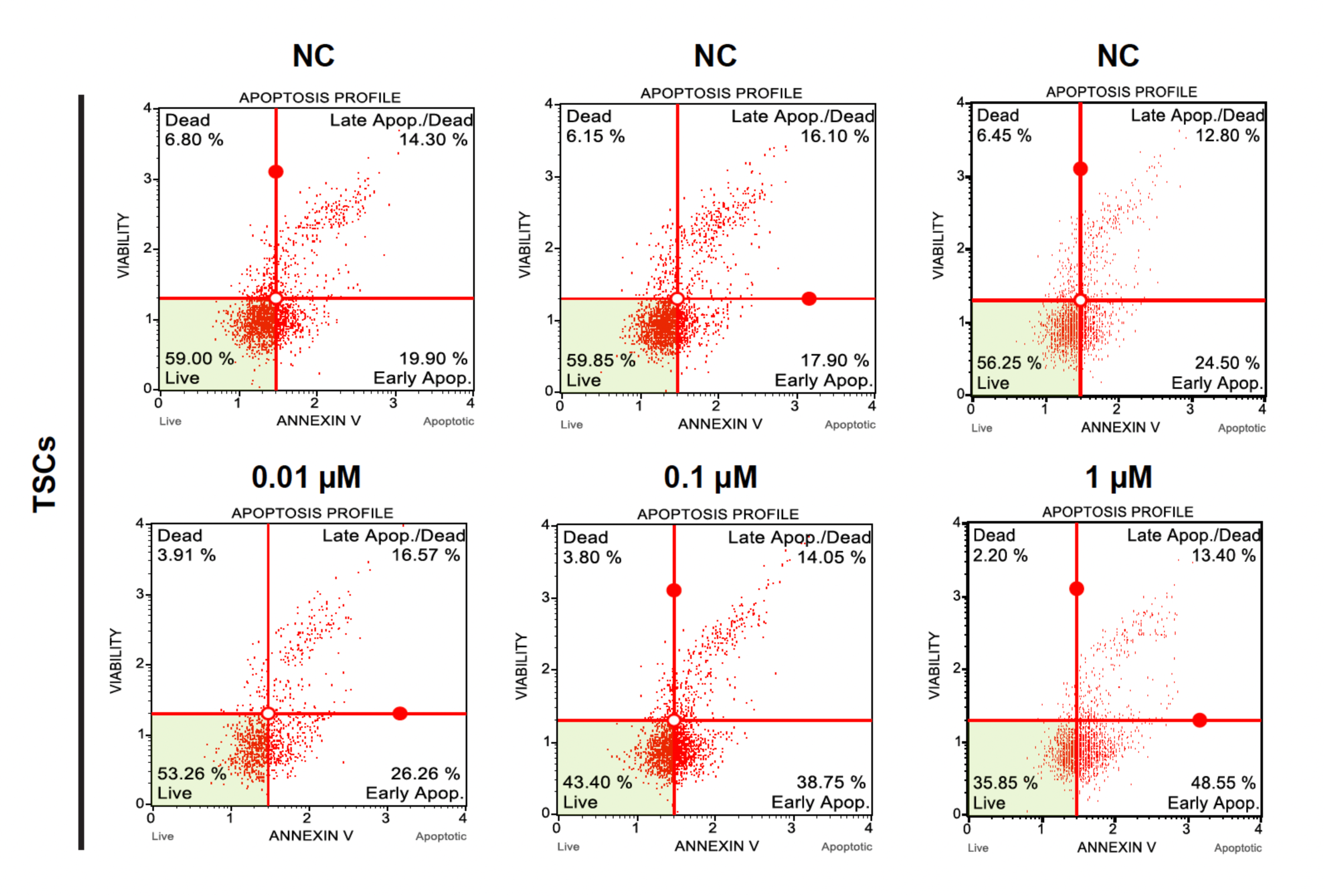
Representative graphs showing the proportion of apoptotic cells among the BPS-treated TSCs and NCs.

**Supplementary Fig.3.**
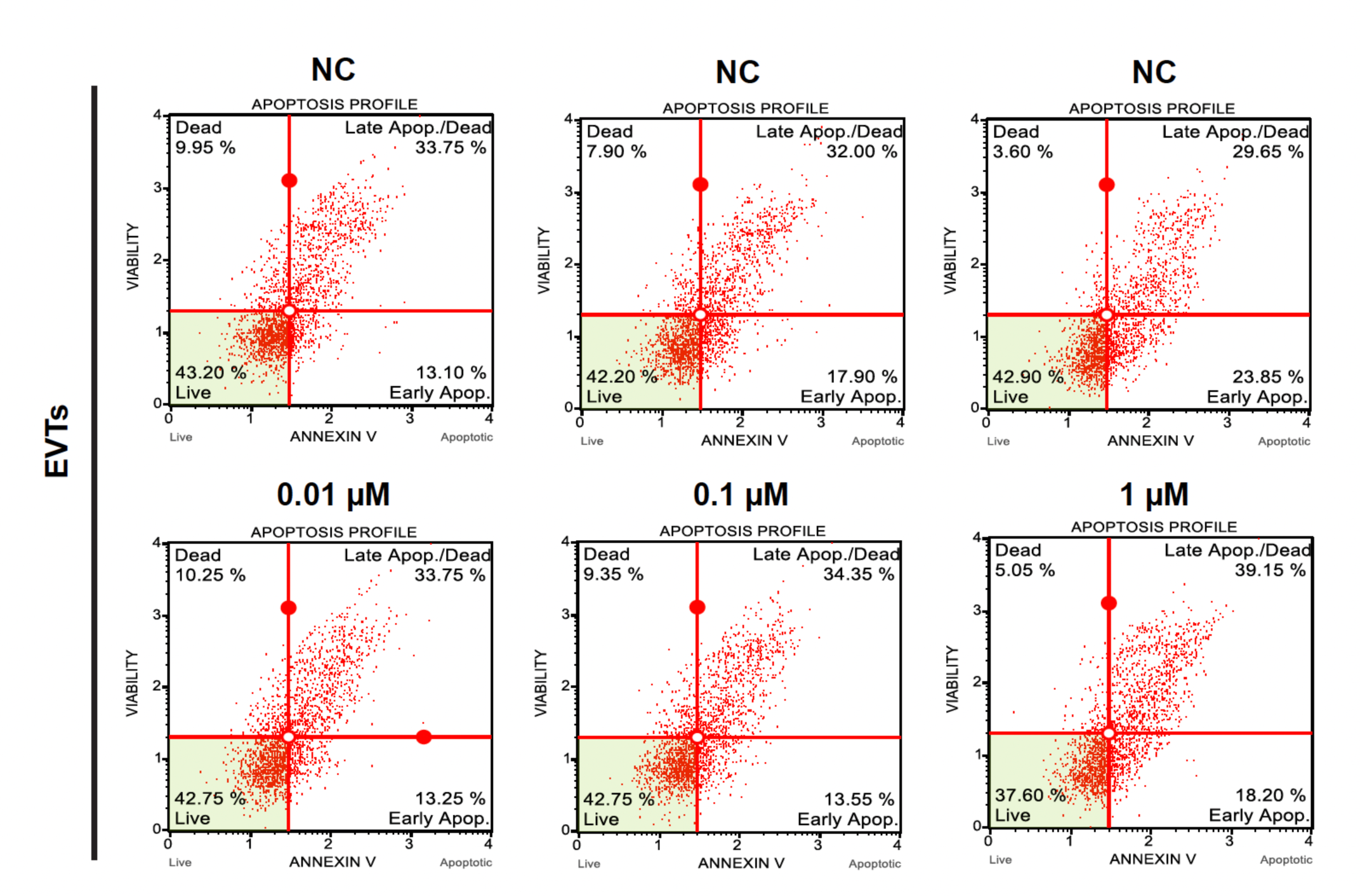
Representative graphs showing the proportion of apoptotic cells among the BPS-treated EVTs and NCs.

**Supplementary Fig.4.**
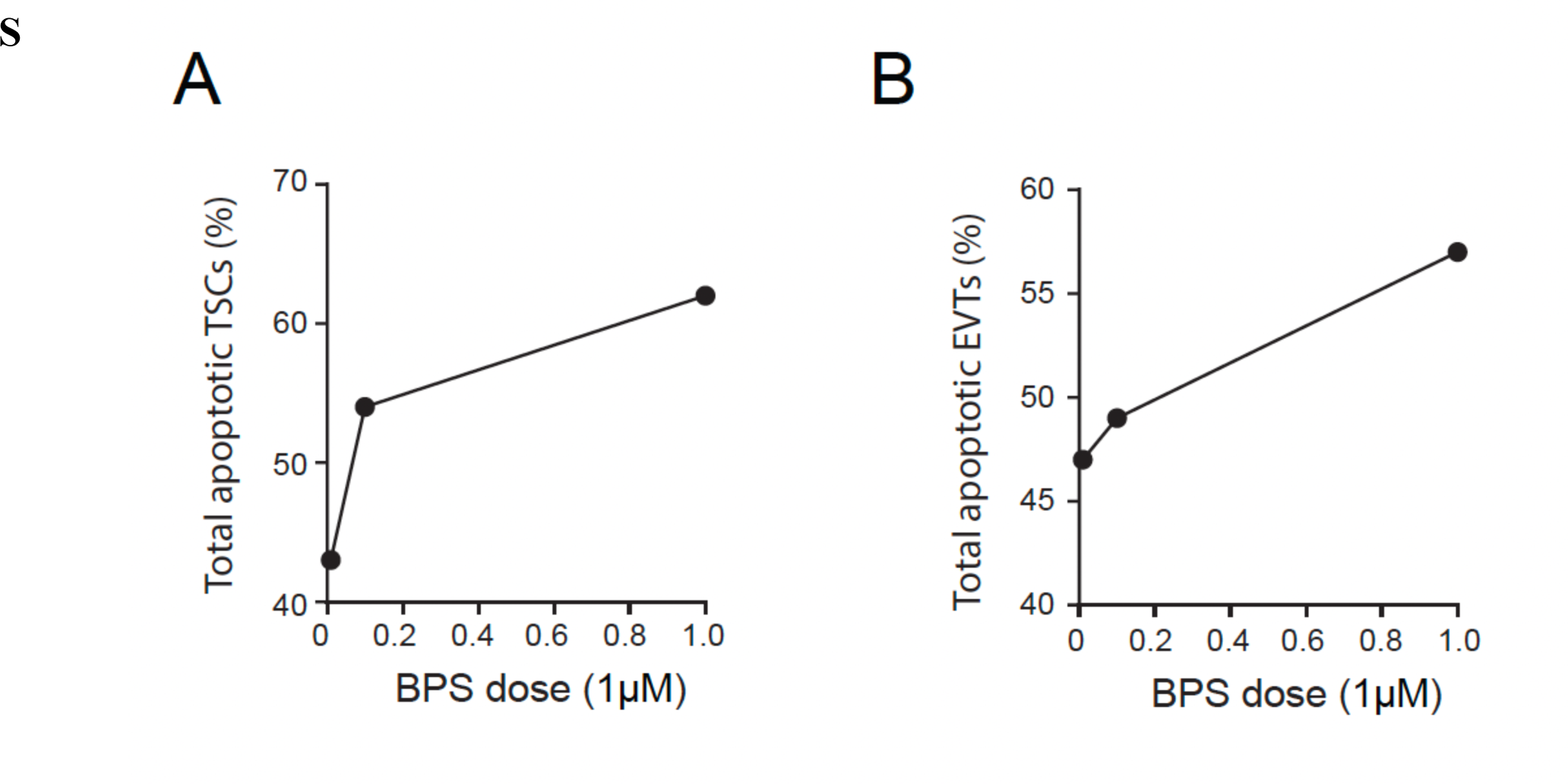
Line graphs showing the dose-dependent relationship between BPS and apoptotic TSCs (A) and EVTs (B)

**Supplementary Table 1.**
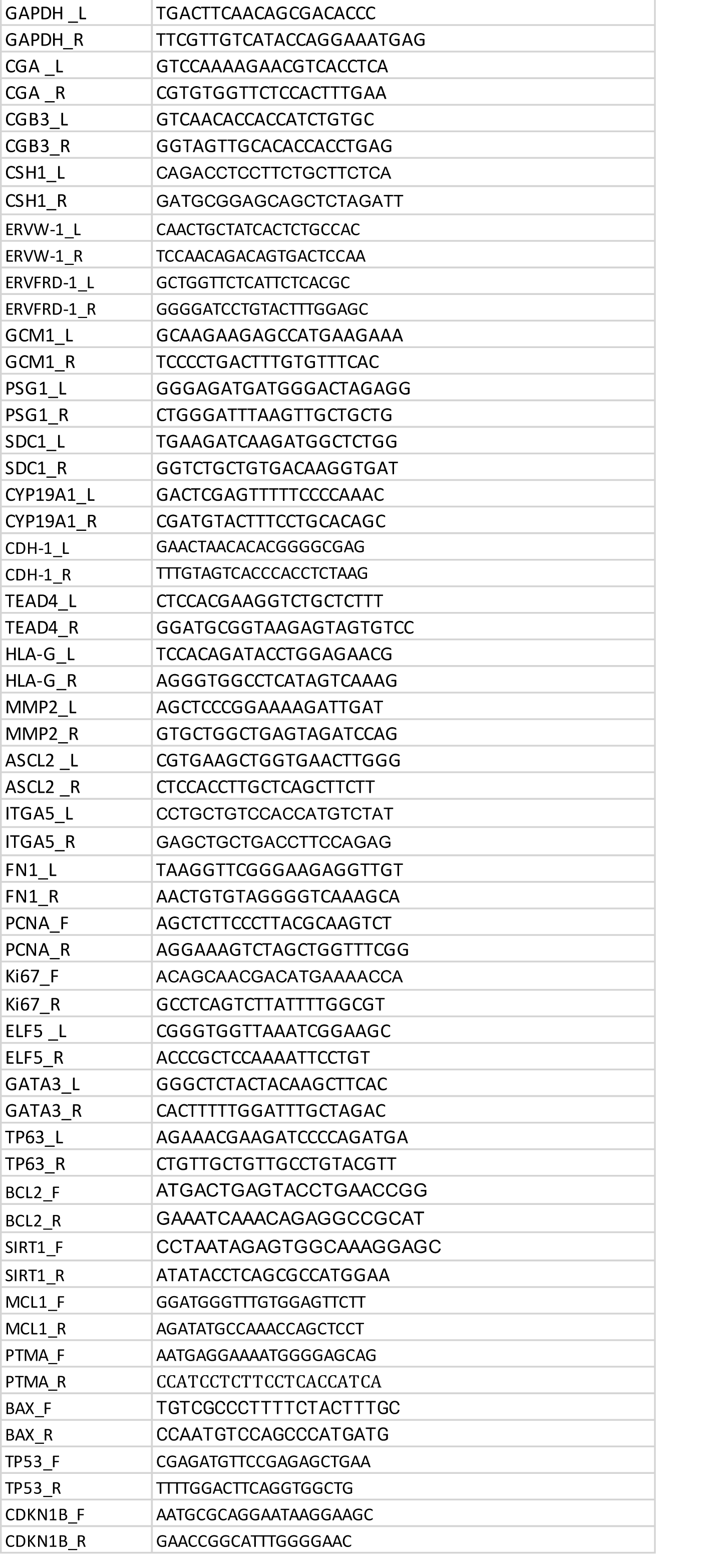
A list of primers ued for quantitative PCR.

